# Impact of prior infection on SARS-CoV-2 antibody responses in vaccinated long-term care facility staff

**DOI:** 10.1101/2022.04.04.487083

**Authors:** Emily N Gallichotte, Mary Nehring, Sophia Stromberg, Michael C Young, Ashley Snell, Josh Daniels, Kristy L Pabilonia, Sue VandeWoude, Nicole Ehrhart, Gregory D Ebel

## Abstract

SARS-CoV-2 emerged in 2019 and has resulted in millions of deaths worldwide. Certain populations are at higher risk for infection, especially staff and residents at long term care facilities (LTCF), due to the congregant living setting, and residents with many comorbidities. Prior to vaccine availability, these populations represented a large fraction of total COVID-19 cases and deaths in the U.S. Due to the high-risk setting and outbreak potential, staff and residents were among the first groups to be vaccinated. To define the impact of prior infection on response to vaccination, we measured antibody responses in a cohort of staff members at a LTCF, many of whom were previously infected by SARS-CoV-2. We found that neutralizing, receptor-binding-domain (RBD) and nucleoprotein (NP) binding antibody levels were significantly higher post-full vaccination course in individuals that were previously infected, and NP antibody levels could discriminate individuals with prior infection from vaccinated individuals. While an anticipated antibody titer increase was observed after vaccine booster dose in naïve individuals, boost response was not observed in individuals with previous COVID-19 infection. We observed a strong relationship between neutralizing antibodies and RBD-binding antibodies post-vaccination across all groups, suggesting RBD-binding antibodies may be used as a correlate of neutralization. One individual with high levels of neutralizing and binding antibodies experienced a breakthrough infection (prior to the introduction of Omicron), demonstrating that the presence of antibodies is not always sufficient for complete protection against infection. These results highlight that history of COVID-19 exposure significantly increases SARS-CoV-2 antibody responses following vaccination.

**Importance:** Long-term care facilities (LTCFs) have been disproportionately impacted by COVID-19, due to their communal nature, high-risk profile of residents and vulnerability to respiratory pathogens. In this study, we analyzed the role of prior natural immunity to SARS-CoV-2 on post-vaccination antibody responses. The LTCF in our cohort experienced a large outbreak with almost 40% of staff becoming infected. We found that individuals that were infected prior to vaccination, had higher levels of neutralizing and binding antibodies post-vaccination. Importantly, the second vaccine dose significantly boosted antibody levels in those that were immunologically naïve prior to vaccination, but not those that had prior immunity. Regardless of pre-vaccination immune status, levels of binding and neutralizing antibodies were highly correlated. The presence of NP-binding antibodies can be used to identify individuals that were previously infected when pre-vaccination immune status is not known. Our results reveal that vaccination antibody responses differ depending on prior natural immunity.

## Introduction

SARS-CoV-2, the virus responsible for COVID-19, has resulted in over 400 million infections worldwide, with 78 million occurring in the United States [1]. Infections in staff and residents of long-term care facilities (LTCFs) account for ~2 million of those infections, and represent 16% of all COVID-19 deaths in the US [2]. LTCFs are high-risk environments due to their congregant living setting, and residents that often have multiple co-morbidities including diabetes, lung and heart disease [3–5]. Because of this, LTCFs have been at the forefront in surveillance testing to detect infections in staff and residents before they spread and cause outbreaks [6, 7]. Additionally, staff and residents at LTCFs were prioritized as one of the first groups to receive vaccines once available, and as of February 2022, over 80% of staff and residents are fully vaccinated nationally [2].

Due to the high number of cases in LTCFs prior to vaccines and other preventative measures, many staff and residents became infected during 2020 and 2021, with some facilities reporting infection and seroprevalence rates as high as 40% [8–11]. Therefore, there are two immunologically distinct populations of individuals receiving vaccines; those that are naïve, with no evidence of a prior infection (seronegative), and those that are pre-immune with either a documented prior infection, or serological evidence of prior infection (seropositive). Early work examined the role of pre-existing immunity on the level of binding antibodies up to four weeks following a single dose of an mRNA vaccine (both Pfizer and Moderna) and found that levels were higher in those that were seropositive [12]. Additional work has evaluated longer term responses after two vaccine doses, and similarly found that those with prior infections generated higher levels of binding antibodies [13–15]. Most of these studies do not measure polyclonal antibody neutralization of live SARS-CoV-2 virus, and instead use pseudotyped virus, or receptor blocking assays as surrogates of true neutralization.

Staff at a local long-term care facility (LTCF), in parallel with their weekly SARS-CoV-2 nasal surveillance qPCR testing, provided blood samples for antibody analyses [8]. This facility experienced a SARS-CoV-2 outbreak in September 2020 prior to vaccine availability, resulting in infection and seroconversion of almost 35% of the staff members [8]. In January 2021, a Pfizer vaccine clinic was provided at their workplace, with the second dose provided three weeks later in early February. Vaccines were not required at this time, though vaccination is now required with rare exceptions [16]. As of January 30, 2022, 96% of staff and 97% of residents at this facility were fully vaccinated, slightly higher than Colorado statewide averages (92% and 93% of staff and residents respectively) [2]. We collected and analyzed sera from staff at this facility from August to December 2020 [8]. We found that during an outbreak at the facility, many staff (~30%) became infected and subsequently seroconverted, generating neutralizing, spike- and RBD-binding antibodies. Here we report seroantibody levels detected in samples collected from February through September 2021 to examine humoral immune response duration. We characterized antibody neutralization, binding to receptor-binding domain (RBD, contained within the spike protein component of the vaccine), and nucleocapsid (NP, not present within the mRNA vaccines). We found that individuals with a prior SARS-CoV-2 infection, have higher post-vaccination neutralizing, RBD and NP binding antibodies than those that were seronegative prior to vaccination, and individuals that were never infected with SARS-CoV-2 did not harbor anti-NP seroreactivity.

## Material and Methods

### Human specimens

This study was approved by the Colorado State University Institutional Review Board under protocol number 20-10057H. Participation in providing blood samples was voluntary. Participants were consented and enrolled and informed of test results. Staff represented a range of job classifications, including those in direct patient care roles (e.g. nurses) and non-direct patient care roles (e.g. administrative).

### Serum collection

Whole blood was collected in BD Vacutainer blood collection tubes and allowed to clot at room temperature for at least 30 minutes. Tubes were spun at 1,300xg for 10 minutes to separate sera from the blood clot. Sera was aliquoted, heat inactivated at 56°C for 30 minutes, and stored at 4°C.

### Viruses and cells

Vero cells (ATCC-81) were maintained in DMEM with 10% fetal bovine serum (FBS), and 1% antibiotic/antimycotic at 37°C and 5% CO_2_. SARS-CoV-2 virus (2019-nCoV/USA-WA1/2020 strain) was used to infect Vero cells for 3 days, supernatant was harvested, centrifuged at maximum speed for 10 minutes to pellet cell debris, aliquoted into single-use aliquots, and stored at −80°C until use.

### Neutralization assay

Standard plaque reduction neutralization test (PRNT) was performed as previously described [8]. Briefly, diluted sera were mixed with virus, incubated for one hour at 37°C, added to a Vero cell monolayer, incubated an additional hour at 37°C, then overlaid with tragacanth media and incubated for two days. Cells were fixed and stained with ethanol and crystal violet, and plaques counted manually.

### RBD and NP ELISA

Binding assays were performed as described previously [8]. Briefly, 96-well plates were coated with SARS-CoV-2 protein (RBD and NP from Sino Biological), blocked with non-fat dried milk, and diluted sera was added. Plates were washed and a secondary anti-human IgG-horseradish peroxidase conjugated secondary antibody was added. Plates were developed and read at 490nm on a spectrophotometer.

### Surveillance qPCR testing

Surveillance testing was performed as previously described [8, 9]. Briefly, nasal swabs were collected, processed, viral RNA extracted, and quantitative reverse transcriptase PCR (qPCR) was performed using the Thermo Fisher Scientific TaqPath COVID-10 combo kit, under U.S. FDA Emergency Use Authorization [17].

## Results

### Neutralizing serum antibodies increase following vaccination regardless of pre-vaccination immune status

By February 2021, neutralizing antibody levels of most individuals increased following one or two doses of vaccine. By mid-March 2021, almost all participating staff had detectable neutralizing antibody levels (**Figure 1A**). Based on prior surveillance testing and antibody analyses [8], we stratified individuals based on their December 2020 immune status as either seropositive (immune), seronegative (naïve), or unknown. On average, immune individuals had higher levels of neutralizing antibodies than those that were seronegative (**Figure 1B**). Not all individuals within this cohort were vaccinated, and in these individuals, neutralizing antibodies detected resulted from natural infection and not vaccination. We next focused on vaccinated individuals, and analyzed neutralizing antibody response based on time post first vaccine dose (ranging from December 2020 to August 2021). When analyzed by days post first vaccine dose, we demonstrated a rapid increase of neutralizing antibody levels (**Figure 1C**). When stratified by pre-vaccination immune status, those who were previously infected had higher levels of neutralizing antibodies post-vaccination compared to individuals that were seronegative prior to vaccination (**Figure 1D**).

**Figure 1.**
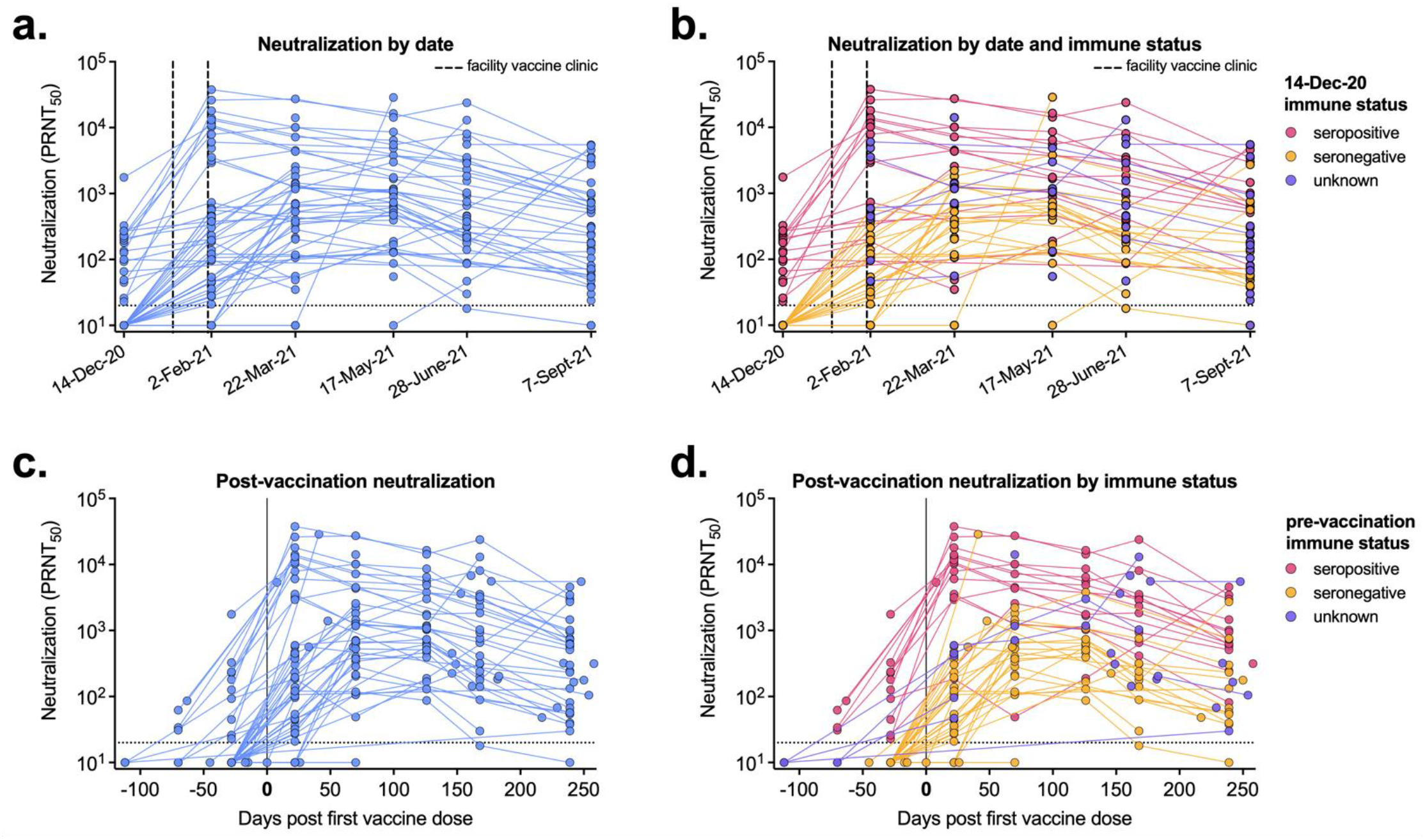
Post-vaccination serum neutralizing levels vary by prior infection. Neutralization titers (PRNT_50_) for each sera sample are shown based on **A, B)** blood collection date or **C, D)** by days post first vaccine dose. **B, D)** Serum samples are labeled based on their pre-vaccination immune status; seropositive (red), seronegative (green), unknown immune status (gray). Dashed line represents the limit of detection (20). Samples without neutralization detected are plotted at half the limit of detection (10).

### Receptor binding domain (RBD) and nucleoprotein (NP) antibody levels after vaccination

We next measured RBD and NP antibody binding levels following vaccination in our cohort participants. RBD levels reached their maximum level in all individuals by day 70 post-vaccination and gradually decreased over the next 6 months (**Figure 2A**). Seropositive individuals had slightly higher RBD absorbance values than those that were immunologically naïve prior to vaccination, though this enhancement was not as marked as neutralizing antibody levels (**Figure 2B**). Participants in our cohort received either the Pfizer or Moderna mRNA vaccines, which encode the viral spike protein (which contains the RBD). Therefore, as expected, only participants with NP-reactive antibodies (**Figure 2C**) were previously infected with SARS-CoV-2 (**Figure 2D**).

**Figure 2.**
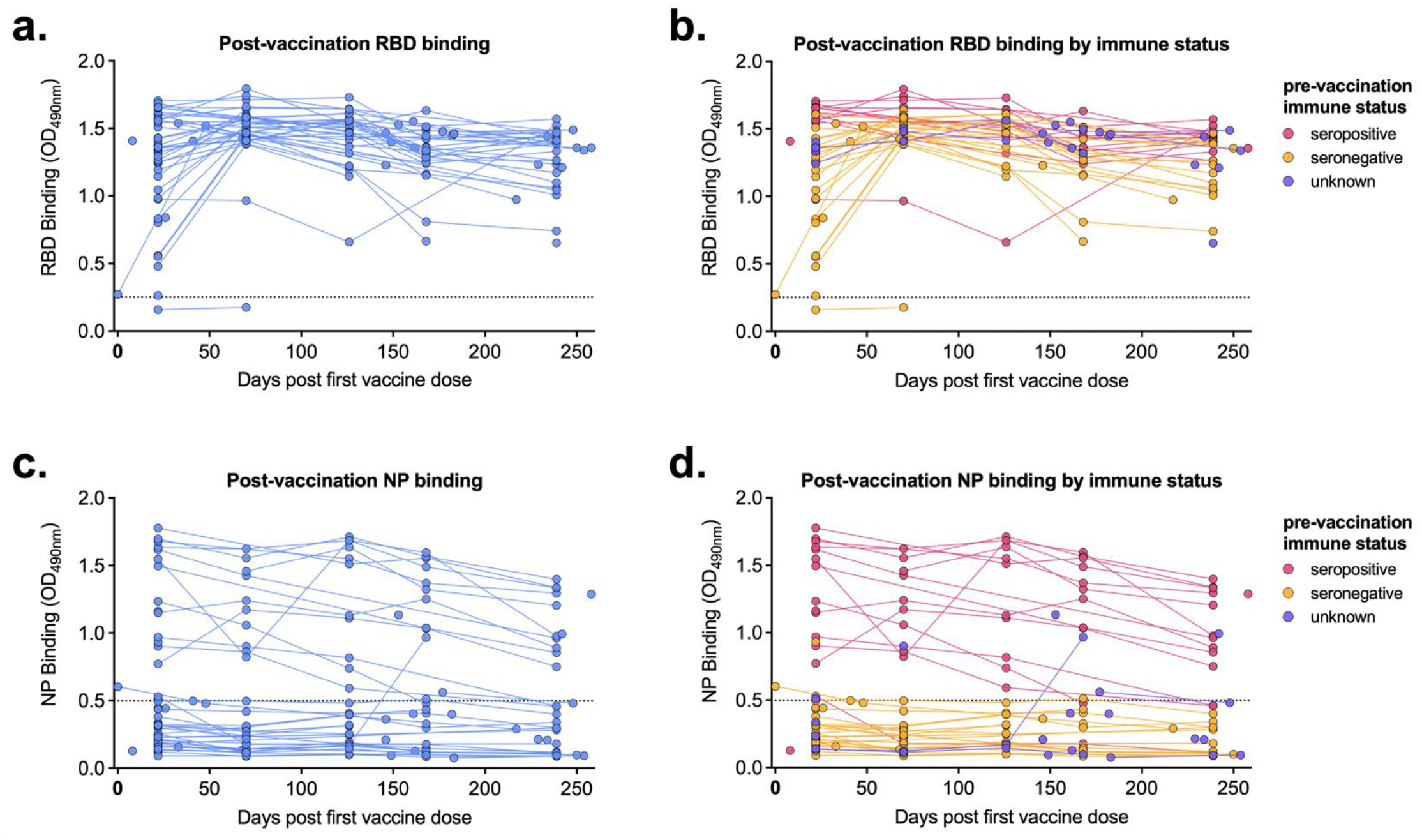
Post-vaccination RBD and NP-binding levels are higher in previously infected individuals. **A, B)** Receptor binding domain (RBD) and **C, D)** nucleoprotein (NP) binding levels for each sera sample are shown by days post first vaccine dose. **B, D)** Serum samples are labeled based on their pre-vaccination immune status; seropositive (red), seronegative (green), unknown immune status (gray). Dashed line represents background level for each assay.

### Post-vaccination antibody levels are higher in pre-immune individuals

When compiling all samples collected post-vaccination (including those after only the first dose), we saw that seropositive pre-vaccination individuals had significantly higher (p<0.0001) levels of neutralizing and RBD binding antibodies compared to seronegative individuals (**Figure 3A & B**). Since there is no nucleoprotein component in the vaccine, it is not surprising that only individuals that experienced a SARS-CoV-2 infection prior to vaccination had detectable NP antibodies (**Figure 3C**). From these results, we can presume that individuals with unknown pre-vaccination immune status (dark gray) with detectable NP-binding antibodies (a single sample from three participants), experienced a SARS-CoV-2 infection prior to vaccination.

**Figure 3.**
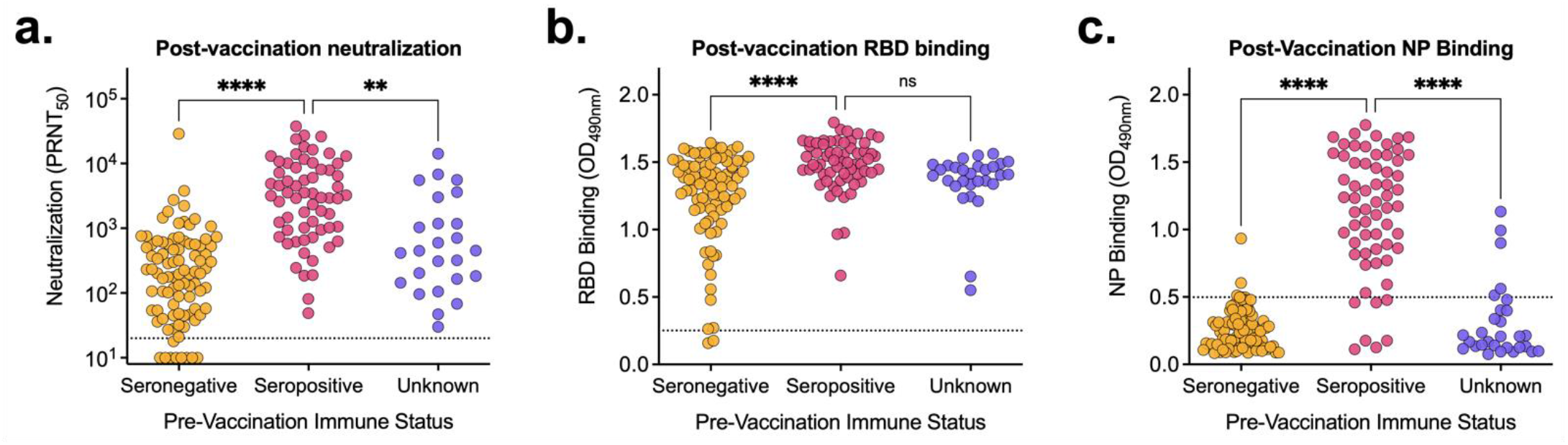
Pre-vaccination immune status impacts post-vaccination antibody levels. All post-vaccination **A)** neutralization titers, **B)** RBD binding and **C)** NP binding values were aggregated and stratified based on pre-vaccination immune status; seropositive (red), seronegative (green), unknown immune status (gray). **A)** dashed lines represent the limit of detection. Samples without neutralization detected are plotted at half the limit of detection (10). **B, C)** Dashed line represents the background level for each assay. **p<0.01, ****p<0.0001, by Tukey’s multiple comparisons one-way ANOVA.

### Impact of second vaccine dose on antibody levels is dependent on pre-vaccination immune status

A subset of the cohort with known serostatus prior to vaccination provided blood samples following both their first and second vaccine doses. We compared levels of neutralizing, RBD binding, and NP binding across these two time points and cohorts and looked at relative changes in antibody levels (**Figure 4**). In immunologically naïve individuals prior to vaccination, neutralizing and RBD binding levels significantly increased between first and second doses (p<0.001) (**Figure 4A & B**). Importantly, some individuals did not have detectable neutralizing antibodies until after their second dose. In contrast, in previously infected individuals, neutralizing and RBD binding antibody levels did not significantly increase following their second dose (**Figure 4A & B**). Additionally, vaccination did not alter NP antibody binding levels regardless of pre-vaccination immune status (**Figure 4C**). In seronegative individuals, following the second vaccine dose, neutralizing and RBD binding levels increased significantly (average of 17-fold and 1.5-fold respectively) (**Figure 4D & E**). Conversely, in pre-vaccination seropositive individuals, on average, neutralizing, RBD binding, and NP binding antibody levels did not change following the second vaccine dose (0.7-, 1- and 0.9-fold changes respectively) (**Figure 4D, E & F**).

**Figure 4.**
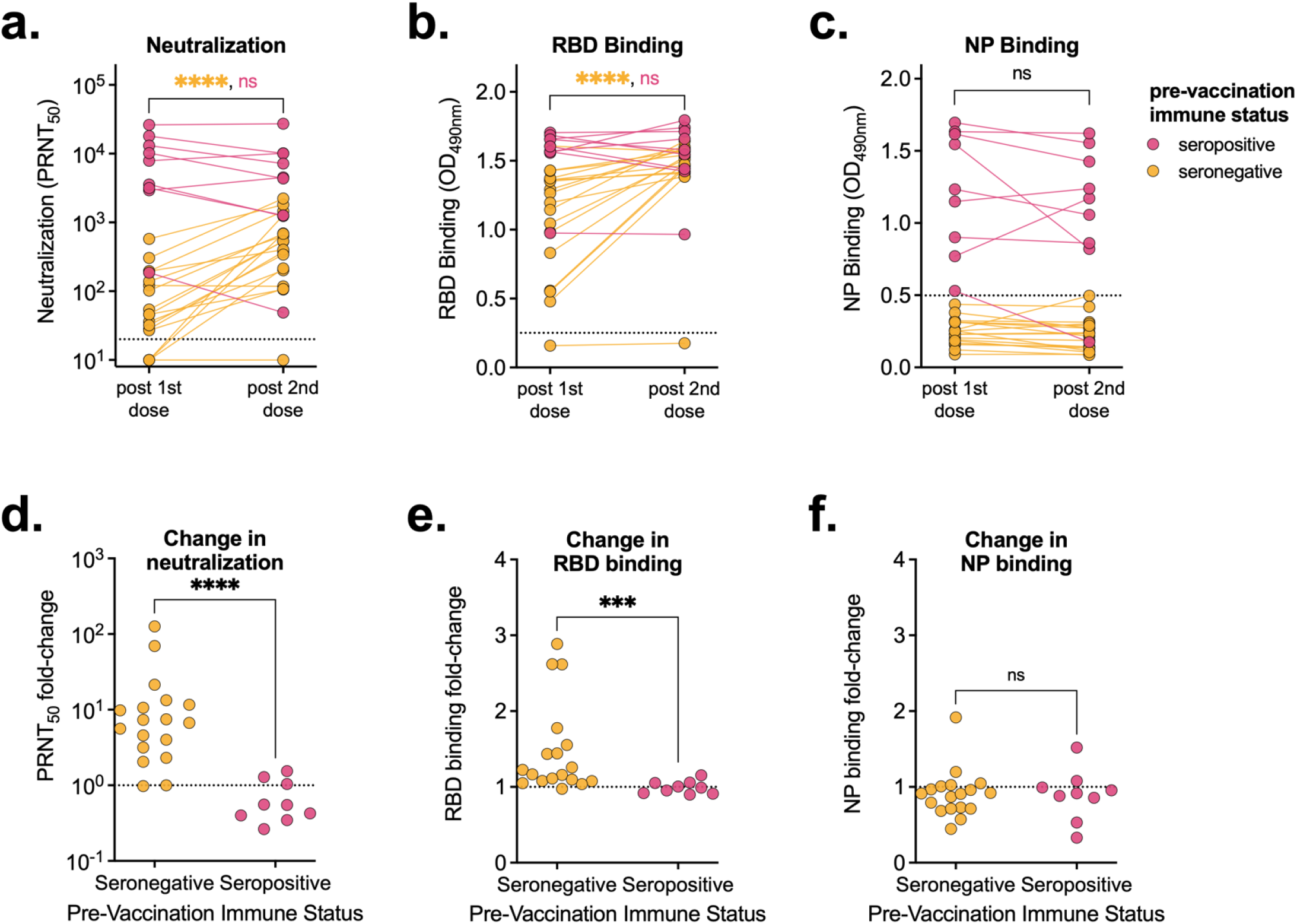
Second vaccine dose only increases antibody levels in seronegative individuals. **A)** Neutralization titers, **B)** RBD binding and **C)** NP binding of serum from individuals following their first and second vaccine doses (3 weeks after the first dose, and 7 weeks after the second dose), stratified by pre-vaccination immune status. Fold change between **D)** neutralization, **E)** RBD binding and **F)** NP binding relative to levels following their first vaccine dose. **A)** dashed lines represent the limit of detection. Samples without neutralization detected are plotted at half the limit of detection (10). **B, C)** Dashed line represents the background level for each assay. ***p<0.001, ****p<0.0001 by Mann-Whitney test.

### Relationship between neutralizing and binding antibodies in vaccinated individuals

We next compared the relationship between neutralizing and binding (both RBD and NP) antibodies in vaccinated individuals (including samples collected after just the first dose), stratified by pre-vaccination immune status. We saw a strong relationship (r > 0.75) between neutralizing titer and RBD binding antibody absorbance regardless of immune status **(Figure 5A**). Because NP antibodies are only found in individuals that experienced a natural SARS-CoV-2 infection, the relationships with NP antibodies (both PRNT_50_ vs NP and RBD vs NP) were poorly correlated (r < 0.45) in post-vaccination sera samples (**Figure 4B & C**).

**Figure 5.**
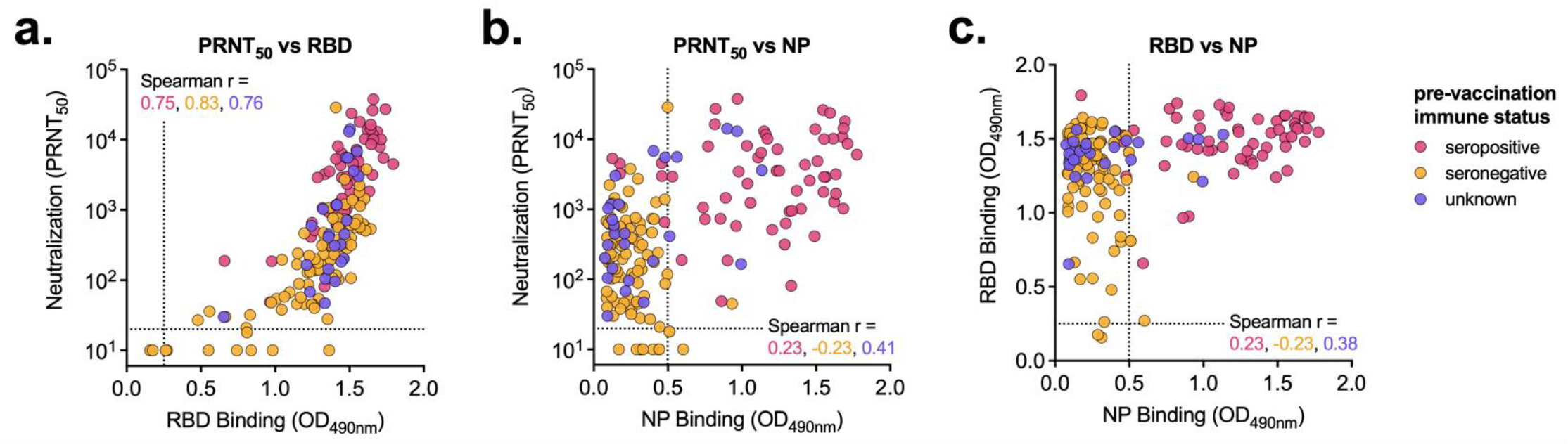
Vaccine-elicited antibody levels are similarly correlated regardless of immune status. Post-vaccination sera samples were compared by **A)** neutralization vs RBD binding, **B)** neutralization vs NP binding and **C)** RBD binding vs NP binding. Serum samples are labeled based on their pre-vaccination immune status; seropositive (red), seronegative (green), unknown immune status (gray). Neutralization dashed line represents the limit of detection. Samples without neutralization detected are plotted at half the limit of detection. RBD and NP dashed line represents the background level for each assay. Spearman r values for each group (seropositive, seronegative, and unknown) are noted.

### Breakthrough infection in a vaccinated individual with high levels of antibodies

One individual in the cohort experienced a breakthrough infection. This individual was seronegative prior to vaccination (no evidence of neutralizing antibodies, nor had they ever tested positive during weekly surveillance testing) and received both vaccine doses in early 2021. In May 2021, this individual experienced an asymptomatic acute breakthrough infection prior to the introduction of the Omicron variant (**Figure 6A**). There was no evidence that antibody levels had waned prior to infection (**Figure 6B & C)**. Neutralizing antibody levels rapidly increased following infection (**Figure 6B**) whereas their RBD binding antibodies did not (**Figure 6C**). Detection of anti-NP antibodies confirmed the breakthrough infection (**Figure 6D**).

**Figure 6.**
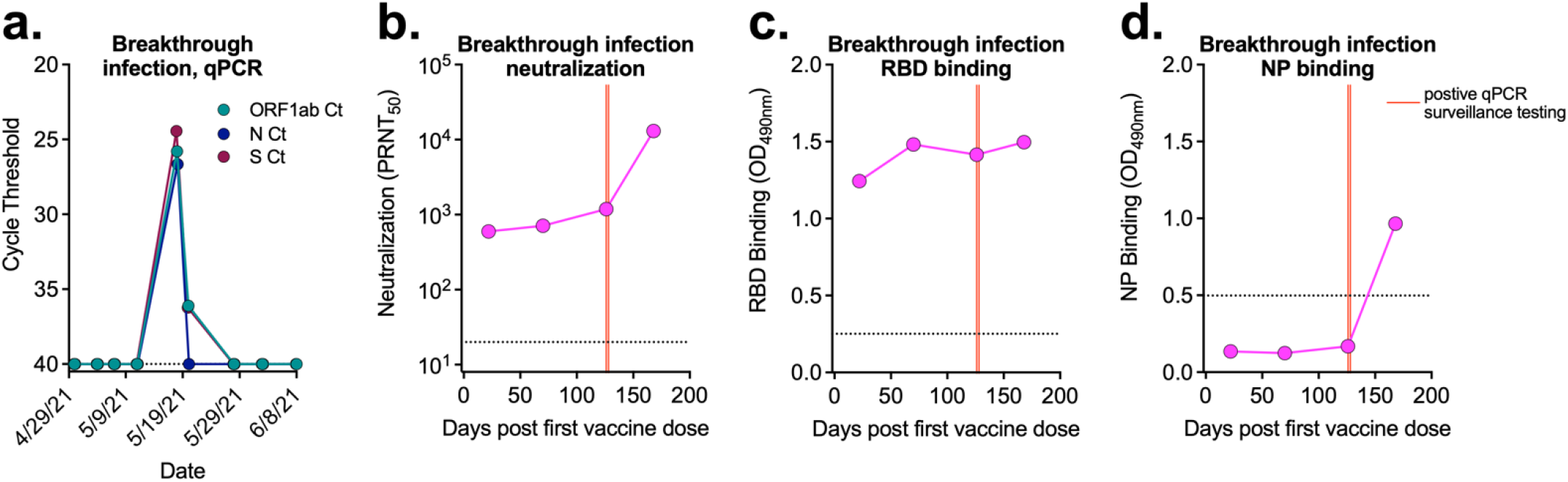
Breakthrough infection in a vaccinated individual. **A)** qPCR surveillance testing for three viral targets from the individual with a breakthrough infection that was seronegative prior to vaccination. **B)** Neutralization, **C)** RBD binding and **D)** NP binding post-vaccination, prior to and post breakthrough infection. Red lines correspond to dates of positive qPCR surveillance testing. **A, B)** Dashed line represents limit of detection. **C, D)** Dashed line represents the background level for each assay.

## Discussion

Early following SARS-CoV-2 vaccine approval, it was unclear if both doses of the mRNA vaccine would be necessary for individuals that had previously been infected, to achieve full protection [18]. It was predicted the first dose would boost humoral immunity acquired from a natural infection. Multiple studies have demonstrated that in previously infected, seropositive individuals, a single vaccine dose is sufficient to generate robust immune responses (both humoral and cellular), often to levels higher than in naïve individuals that received two vaccine doses [19–21]. Our data confirm that individuals with a prior infection generate a robust, neutralizing antibody response that is not further increased upon a second dose. These results have led to calls for a single-dose vaccine regimen in previously infected individuals to stretch vaccine supplies, improve worldwide vaccine access, and increase vaccine uptake among hesitant COVID-19 survivors [22–25].

Conversely, in seronegative individuals, antibody levels significantly increased following a second vaccine dose [19–21]. Three individuals in our cohort did not generate neutralizing antibodies until after the second vaccine dose, and one individual never seroconverted following vaccination. It is therefore critical that individuals without prior infection receive the full vaccination course to ensure maximum immune response [26].

Neutralizing and binding antibody levels are being developed as correlates of protection, as they are highly correlated with vaccine efficacy across diverse cohorts and vaccine platforms [27–29]. There are reports describing breakthrough infections post-vaccination, likely due to reduced/waning antibody levels and timing post-vaccination [30–33]. The breakthrough infection that occurred in our cohort, was in an individual with high neutralizing antibody levels similar to other recent reports [34, 35]. These data suggest that while antibody levels may be broadly predictive of vaccine efficacy, they are not sufficient as a singular correlate of protection in all individuals.

Our work, along with others [36–38], describes the use of nucleoprotein antibody detection as a tool to identify natural infection using serum collected post-vaccination. This assay could be used to further define and refine correlates of protection, or generate a better predictor of breakthrough risk, by stratifying post-vaccination serum into those that had and had not been previously infected. Importantly, this strategy is only effective in individuals that received a vaccine without a nucleocapsid component (Pfizer, Moderna, etc.) as opposed to inactivated whole virus vaccines (or other similar vaccine platforms) containing nucleocapsid, such as Sinovac.

There remain many unknowns regarding the immune response following COVID-19 infection, vaccination, booster, and breakthrough [39–41]. Boosters, which have been widely accessible in the U.S., combat waning immunity by boosting pre-existing adaptive immunity (both humoral and cellular), furthering protection against severe disease [42]. There is relatively high booster uptake among staff and residents of LTCFs in Colorado (76% and 40% of residents and staff with boosters, respectively), with slightly higher rates in the facility described in this paper (80% residents, 44% staff) [2]. Despite high vaccination and booster rates, the Omicron variant seems to efficiently evade vaccine-elicited immunity [43, 44]. These results suggest additional boosters, and variant-specific boosters might be required to maintain long-term immunity against SARS-CoV-2 [45].

## Acknowledgements

This work was supported with funds from the Boettcher Foundation and funds donated by the Colorado State University College of Health and Human Services, Veterinary Medicine and Biomedical Sciences, Natural Sciences and Walter Scott, Jr. College of Engineering, the Colorado State University Columbine Health Systems Center for Health Aging, and the CSU One Health Institute. We thank the members of the CSU Veterinary Diagnostic Lab for their assistance with surveillance testing and diagnostic support. Additionally, we thank the ongoing support and participation of the LTCF staff who participated in the study.

## References

1. Times, T.N.Y. Coronavirus in the U.S.: Latest Map and Case Count. 2022 February 16, 2022 [cited 2022 February 16]; Available from: https://www.nytimes.com/interactive/2021/us/covid-cases.html.

2. Services, C.f.M.M. COVID-19 Nursing Home Data. 2022 February 10, 2022 [cited 2022 February 16]; Available from: https://data.cms.gov/covid-19/covid-19-nursing-home-data.

3. Ochieng, N., et al., Factors Associated With COVID-19 Cases and Deaths in Long-Term Care Facilities: Findings from a Literature Review. 2021, Kaiser Family Foundation: kff.org.

4. Tang, O., et al., Outcomes of Nursing Home COVID-19 Patients by Initial Symptoms and Comorbidity: Results of Universal Testing of 1970 Residents. J Am Med Dir Assoc, 2020. 21(12): p. 1767–1773 e1.

5. Ouslander, J.G. and D.C. Grabowski, COVID-19 in Nursing Homes: Calming the Perfect Storm. J Am Geriatr Soc, 2020. 68(10): p. 2153–2162.

6. Litwin, T., et al., Preventing COVID-19 outbreaks through surveillance testing in healthcare facilities: a modelling study. BMC Infect Dis, 2022. 22(1): p. 105.

7. Smith, D.R.M., et al., Optimizing COVID-19 surveillance in long-term care facilities: a modelling study. BMC Med, 2020. 18(1): p. 386.

8. Gallichotte, E.N., et al., Durable Antibody Responses in Staff at Two Long-Term Care Facilities, during and Post SARS-CoV-2 Outbreaks. Microbiol Spectr, 2021. 9(1): p. e0022421.

9. Gallichotte, E.N., et al., Early Adoption of Longitudinal Surveillance for SARS-CoV-2 among Staff in Long-Term Care Facilities: Prevalence, Virologic and Sequence Analysis. Microbiol Spectr, 2021. 9(3): p. e0100321.

10. Wisniak, A., et al., Association between SARS-CoV-2 Seroprevalence in Nursing Home Staff and Resident COVID-19 Cases and Mortality: A Cross-Sectional Study. Viruses, 2021. 14(1).

11. Tanunliong, G., et al., Persistence of Anti-SARS-CoV-2 Antibodies in Long Term Care Residents Over Seven Months After Two COVID-19 Outbreaks. Front Immunol, 2021. 12: p. 775420.

12. Krammer, F., et al., Antibody Responses in Seropositive Persons after a Single Dose of SARS-CoV-2 mRNA Vaccine. N Engl J Med, 2021. 384(14): p. 1372–1374.

13. Demonbreun, A.R., et al., Comparison of IgG and neutralizing antibody responses after one or two doses of COVID-19 mRNA vaccine in previously infected and uninfected individuals. EClinicalMedicine, 2021. 38: p. 101018.

14. Fraley, E., et al., Humoral immune responses during SARS-CoV-2 mRNA vaccine administration in seropositive and seronegative individuals. BMC Med, 2021. 19(1): p. 169.

15. Moncunill, G., et al., Determinants of early antibody responses to COVID-19 mRNA vaccines in a cohort of exposed and naive healthcare workers. EBioMedicine, 2022. 75: p. 103805.

16. Services, C.f.M.M., Guidance for the Interim Final Rule - Medicare and Medicaid Programs; Omnibus COVID-19 Health Care Staff Vaccination. 2021.

17. Administration, U.S.F.D., TaqPath COVID-19 Combo Kit - Letter of Authorization. 2021.

18. Dolgin, E., Is one vaccine dose enough if you’ve had COVID? What the science says. Nature, 2021. 595(7866): p. 161–162.

19. Ebinger, J.E., et al., Antibody responses to the BNT162b2 mRNA vaccine in individuals previously infected with SARS-CoV-2. Nat Med, 2021. 27(6): p. 981–984.

20. Goel, R.R., et al., Distinct antibody and memory B cell responses in SARS-CoV-2 naive and recovered individuals following mRNA vaccination. Sci Immunol, 2021. 6(58).

21. Stamatatos, L., et al., mRNA vaccination boosts cross-variant neutralizing antibodies elicited by SARS-CoV-2 infection. Science, 2021.

22. Frieman, M., et al., SARS-CoV-2 vaccines for all but a single dose for COVID-19 survivors. EBioMedicine, 2021. 68: p. 103401.

23. Ledford, H., How can countries stretch COVID vaccine supplies? Scientists are divided over dosing strategies. Nature, 2021. 589(7841): p. 182.

24. UK science advisers: publish evidence behind COVID vaccine changes. Nature, 2021. 589(7841): p. 169–170.

25. Wood, S. and K. Schulman, Beyond Politics - Promoting Covid-19 Vaccination in the United States. N Engl J Med, 2021. 384(7): p. e23.

26. Accorsi, E.K., et al., Association Between 3 Doses of mRNA COVID-19 Vaccine and Symptomatic Infection Caused by the SARS-CoV-2 Omicron and Delta Variants. JAMA, 2022. 327(7): p. 639–651.

27. Cromer, D., et al., Neutralising antibody titres as predictors of protection against SARS-CoV-2 variants and the impact of boosting: a meta-analysis. Lancet Microbe, 2022. 3(1): p. e52–e61.

28. Earle, K.A., et al., Evidence for antibody as a protective correlate for COVID-19 vaccines. Vaccine, 2021. 39(32): p. 4423–4428.

29. Khoury, D.S., et al., Neutralizing antibody levels are highly predictive of immune protection from symptomatic SARS-CoV-2 infection. Nat Med, 2021. 27(7): p. 1205–1211.

30. Evans, J.P., et al., Neutralizing antibody responses elicited by SARS-CoV-2 mRNA vaccination wane over time and are boosted by breakthrough infection. Sci Transl Med, 2022: p. eabn8057.

31. Mizrahi, B., et al., Correlation of SARS-CoV-2-breakthrough infections to time-from-vaccine. Nat Commun, 2021. 12(1): p. 6379.

32. Paniskaki, K., et al., Immune Response in Moderate to Critical Breakthrough COVID-19 Infection After mRNA Vaccination. Front Immunol, 2022. 13: p. 816220.

33. Ahmed, S., et al., Post-vaccination antibody titres predict protection against COVID-19 in patients with autoimmune diseases: survival analysis in a prospective cohort. Ann Rheum Dis, 2022.

34. Rovida, F., et al., SARS-CoV-2 vaccine breakthrough infections with the alpha variant are asymptomatic or mildly symptomatic among health care workers. Nat Commun, 2021. 12(1): p. 6032.

35. Terreri, S., et al., Persistent B cell memory after SARS-CoV-2 vaccination is functional during breakthrough infections. Cell Host Microbe, 2022.

36. Assis, R., et al., Distinct SARS-CoV-2 antibody reactivity patterns elicited by natural infection and mRNA vaccination. NPJ Vaccines, 2021. 6(1): p. 132.

37. Fenwick, C., et al., Changes in SARS-CoV-2 Spike versus Nucleoprotein Antibody Responses Impact the Estimates of Infections in Population-Based Seroprevalence Studies. J Virol, 2021. 95(3).

38. van den Hoogen, L.L., et al., Seropositivity to Nucleoprotein to detect mild and asymptomatic SARS-CoV-2 infections: A complementary tool to detect breakthrough infections after COVID-19 vaccination? Vaccine, 2022.

39. Mohamed, K., et al., COVID-19 vaccinations: The unknowns, challenges, and hopes. J Med Virol, 2022. 94(4): p. 1336–1349.

40. Sette, A. and S. Crotty, Pre-existing immunity to SARS-CoV-2: the knowns and unknowns. Nat Rev Immunol, 2020. 20(8): p. 457–458.

41. Prevention, C.f.D.C.a. Science Brief: SARS-CoV-2 Infection-induced and Vaccine-induced Immunity. 2021 October 29, 2021; Available from: https://www.cdc.gov/coronavirus/2019-ncov/science/science-briefs/vaccine-induced-immunity.html.

42. Prevention, C.f.D.C.a., New CDC Studies: COVID-19 Boosters Remain Safe, Continue to Offer High Levels of Protection Against Severe Disease Over Time and During Omicron and Delta Waves. 2022.

43. Liu, L., et al., Striking Antibody Evasion Manifested by the Omicron Variant of SARS-CoV-2. Nature, 2021.

44. Reardon, S., How well can Omicron evade immunity from COVID vaccines? Nature, 2022.

45. Burki, T.K., Omicron variant and booster COVID-19 vaccines. Lancet Respir Med, 2022. 10(2): p. e17.

